# Entomopathogen-host evolution and implications for biopesticide resistance management

**DOI:** 10.1101/2022.10.28.514078

**Authors:** Jeremy P. Roberts, Tobin D. Northfield

## Abstract

Insecticide resistance evolution is becoming increasingly problematic globally. With chemical insecticides, attempts to combat resistance involves developing compounds with novel modes of action, or increasing rates to overcome partial resistance. While pests can develop resistance to pathogens used as biopesticides, these “pesticides” can be subjected to evolutionary selection pressure as well and may be able to adapt countermeasures to overcome pest resistance. Here, we consider two scenarios: 1) a single trait governs an arms race between pest and parasite, and 2) an epidemiological scenario where each, pathogen transmission and virulence, are governed by host and pathogen traits. Considering the single-trait parasite attack scenario, the evolving parasite is able to overcome resistance in the pest population and effectively suppress host population abundance. In this case, overcoming biopesticide resistance may be possible from parasite evolution to resistant hosts. In contrast, when transmission and abundance are allowed to vary independently in an epidemiological model, different pathogen traits promote different types of resistance development in the host – more contagious pathogens promote pathogen-tolerant (low mortality susceptibility) hosts, while less contagious pathogens promote pathogen-resistant (low transmission susceptibility) hosts. Pathogen-tolerant hosts are particularly detrimental to control programs, because they can quickly outcompete wild types by promoting infection in wild type populations. Furthermore, because evolution of pathogen-tolerance in pests can benefit pathogens through increasing infection prevalence, we do not expect pathogen evolution to improve control. Thus, the keys to biopesticide management depend on the virulence-transmission trade-off and whether hosts evolve to better prevent or survive infection.

## Introduction

The evolution of insecticide resistance in pest species is a major concern in many agricultural systems (FAO, 2012). Widespread use of chemical insecticides has led to many instances of resistance development (Bass et al., 2015). In response to these risks, insecticide management programs typically focus on slowing resistance through conserving wild type pests that can mate with resistant mutants and dilute resistance across agricultural landscapes (Carrière et al., 2010). Because selection typically favors eventual insecticide resistance when the insecticides are applied consistently, management has also included development of novel insecticides and in many cases increasing rates where only partial resistance is present (Bourguet et al., 2013). However, due to the potential for chemical insecticides to damage non-target populations, especially with effects on natural enemies of target species (Tabashnik, 1986), efforts have been made to avoid these increases in insecticide application rates.

Modern molecular techniques have led to improved advances in two types of alternatives to chemical insecticides: genetically based pest management tools (Sun et al., 2017), and pathogens used as biopesticides (Lacey et al., 2015). Aided by improved knowledge of pathogens and the ability to detect novel strains, biopesticides are increasingly being deployed as control measures, partially in response to increased organic production, but also as they offer benefits in resistance management strategies (Lacey et al., 2015). However, even these treatments are susceptible to host resistance evolution. A major example is in codling moth, *Cydia pomonella*, where multiple resistant populations have been found in regard to the biopesticide codling moth granulovirus CpGV (Eberle and Jehle, 2006; Lacey et al., 2008; Zichová et al., 2013). As biopesticide use becomes more widespread, many more of them will likely have to deal with resistances in their hosts. The theory informing management of pest resistance to chemically based insecticides is well established (Furlan et al., 2021; Tabashnik, 1986), and recently theory has been developed to guide the battle resistance to genetically based control methods using gene drive technology such as CRISPR (Gomulkiewicz et al., 2021), but theory focused on biopesticides is lacking. Here, we develop theory to inform management of pest resistant to pathogens and parasites used as biopesticides, guided by coevolutionary arms race models and the evaluation of well-established epidemiological models.

In natural systems, organisms evolve in response to selection pressures exhibited by predators and prey (Hairston Jr et al., 2005). Even in biocontrol programs involving insect control agents, host resistance can pose an issue (Tomasetto et al., 2017). While hosts (or prey) may evolve to better escape consumption, their consumers (predators, parasites, pathogens) can maintain viability by evolving ways to overcome host (or prey) defense (Bohannan and Lenski, 2000; Lively and Dybdahl, 2000; Pimentel, 1968). Biopesticides are typically mass produced in isolation from evolving pest populations and distributed in quantities that may reduce rates of pathogen evolution in nature. However, since these biopesticides are living organisms, they may be able to adapt to evolved defenses in the pest population if given the opportunity to evolve themselves. Ultimately, however, the likelihood and details of this pathogen evolution will depend on the magnitude of any fitness costs associated with its resistance (Bohannan and Lenski, 2000), as well as demographic constraints (Gomulkiewicz and Houle, 2009). Here, we develop theory to evaluate how trait costs and selection pressures can shape evolution of pathogens when allowed to adapt to resistant pest populations. We consider two scenarios: 1) where host death is an important part of transmission as is common in many parasites, for example many entomopathogenic nematodes rely on infectious juveniles and their symbiotic bacteria to kill their insect hosts before conditions become right for them to reproduce (Ciche et al., 2006) and 2) where living, but infected hosts can transmit the pathogen to healthy hosts as is commonly assumed in epidemiological models (Anderson and May, 1979). We term the first scenario as the “Resource-consumer model” and consider the evolution of a single host trait that allows hosts to defend themselves against parasites, and a single parasite trait that allows them to overcome host defenses to kill the host and reproduce. For the second scenario, we evaluate epidemiological models that track the number of infected and susceptible hosts within a population, but do not specifically track the number of pathogens in a population. Here, we focus on host evolution by introducing mutant hosts that differ in their ability to survive or prevent an infection at a cost to reproduction and monitor the long-term survivorship of mutant and wild type hosts.

### Model description

When considering infectious diseases, the traits of transmissibility and virulence are typically used to determine the rate of disease spread. Transmissibility quantifies how likely infection passes from one individual to another, while virulence can describe how likely infection is to be fatal. These traits are often linked, as both can be related to the pathogen load within the host: the more the pathogen replicates within a host, the more likely it can spread but also the more likely it can consume too much of the host’s resources (Antia et al., 1994). Since a dead host would often no longer be suitable habitat for a pathogen, it is faced with a trade-off to balance its virulence with its transmissibility.

If the pathogen’s life cycle is such that host death is necessary for its transmission to occur, as is the case for some entomopathogens, nematodes and others, then virulence and transmission would be completely linked, for example where transmission is a function of virulence. This lends itself to a predator-prey or resource-consumer type modelling approach, where a single consumption efficiency trait effects both the resource’s depletion and growth of the consumer. For comparison, we also consider an epidemiological scenario where pathogens are able to be spread from living infected hosts. This modeling framework follows a susceptible-infected approach developed by Anderson and May (1979).

Of course, for insect pathogens it may not always be clear which framework is most appropriate for a particular insect. For example, transmission may occur between living insects when the densities are high, but primarily occur once the host dies, for example, through a sporulation event (Steinkraus, 2006; Toledo et al., 2007). Thus, we present two scenarios with the understanding that many systems may fall somewhere in between, depending on the rate of transmission from alive versus dead insects. Specifically, we evaluate each of these scenarios to see how biopesticide evolution would be expected to impact pest control and resistance management.

### Resource-consumer model

Here we consider a system where host death is an important part of transmission, where for example the dead host releases infectious spores or juveniles (Ciche et al., 2006; Rohrmann, 2019; Steinkraus, 2006). Since host mortality is a prerequisite of spread (population growth), the system can be thought of as similar to parasitoid and predatory systems and a resource-consumer modeling approach may be appropriate. Models describing these scenarios have been developed for predator-prey interactions (Abrams, 2000; Tien and Ellner, 2012). Here, we adapt one such model (Northfield and Ives, 2013) to describe host-parasite interactions. The approach tracks the two populations of host and parasite. This model incorporates species interaction through population dynamics and quantitative genetics approaches. This species interaction describes the effects of a population’s density on the growth rate of the other. This can be thought of as the attack rate of the parasite on the host, so greater density of either population increases the attack rate for the other, and this would increase parasite growth and decrease host growth. This approach assumes no influence of spatial dynamics on the system, with random mating within each population.

Each population also has a trait that affects the relationship with the other species. In the host, this can be thought of as defensive trait (*v*) that reduces the success of parasite attacks, and in parasite as a trait (*b*) that describes the impacts of these anti-parasite defenses, reducing the success of parasite attack. Thus, the host benefits from having large defensive values (*v*), and the parasite benefits from reduced efficacy of these defenses (*b*). Each trait therefore modifies the successful attack rate, and has an associated trait cost to reproduction rate, which increases with greater trait investment. The traits change over time, dependent on a constant additive genetic variance and the selection pressure (Abrams et al., 1993). This constant variance approach has limitations but should be appropriate for the relatively short time scales of resistance evolution, assuming relatively small values for the genetic variance (*V*_*N*_, *V*_*P*_ << 1) (Abrams, 2001). This genetic variance can be set to 0, to prevent a population from evolving. Here, population of host *N*_*t*_ and parasite *P*_*t*_ at time *t* are described as:

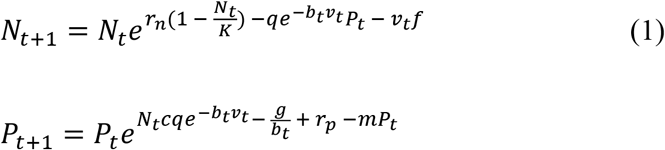

Traits of host *v*_*t*_ and parasite *b*_*t*_ at time *t* are described as follows:

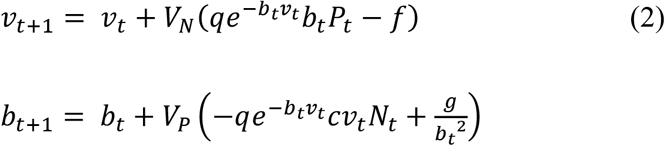

Here, *r*_*n*_ and *r*_*p*_ represent the host and parasite reproduction rates, respectively, while *q* represents the parasitism rate, *m* represents the parasite mortality rate, *c* represents a measure of parasitism efficiency, and *K* represents the host population’s carrying capacity. Further, *f* and *g* represent the degree of trait cost scaling for the host and parasite respectively, and *V*_*N*_ and *V*_*P*_ the genetic variances. The parameter *b*_*t*_ represents the susceptibility of the parasite to host defense, for example the ineffectiveness of the immune response. However, it is often more intuitive to have trait values that are monotonically increasing with investment, so we present in the figures 1-*b*_*t*_ to represent the ability of the parasite to overcome the host’s defense. This parameterization allows for mathematical simplicity while allowing for intuitive interpretation.

**Figure 1:**
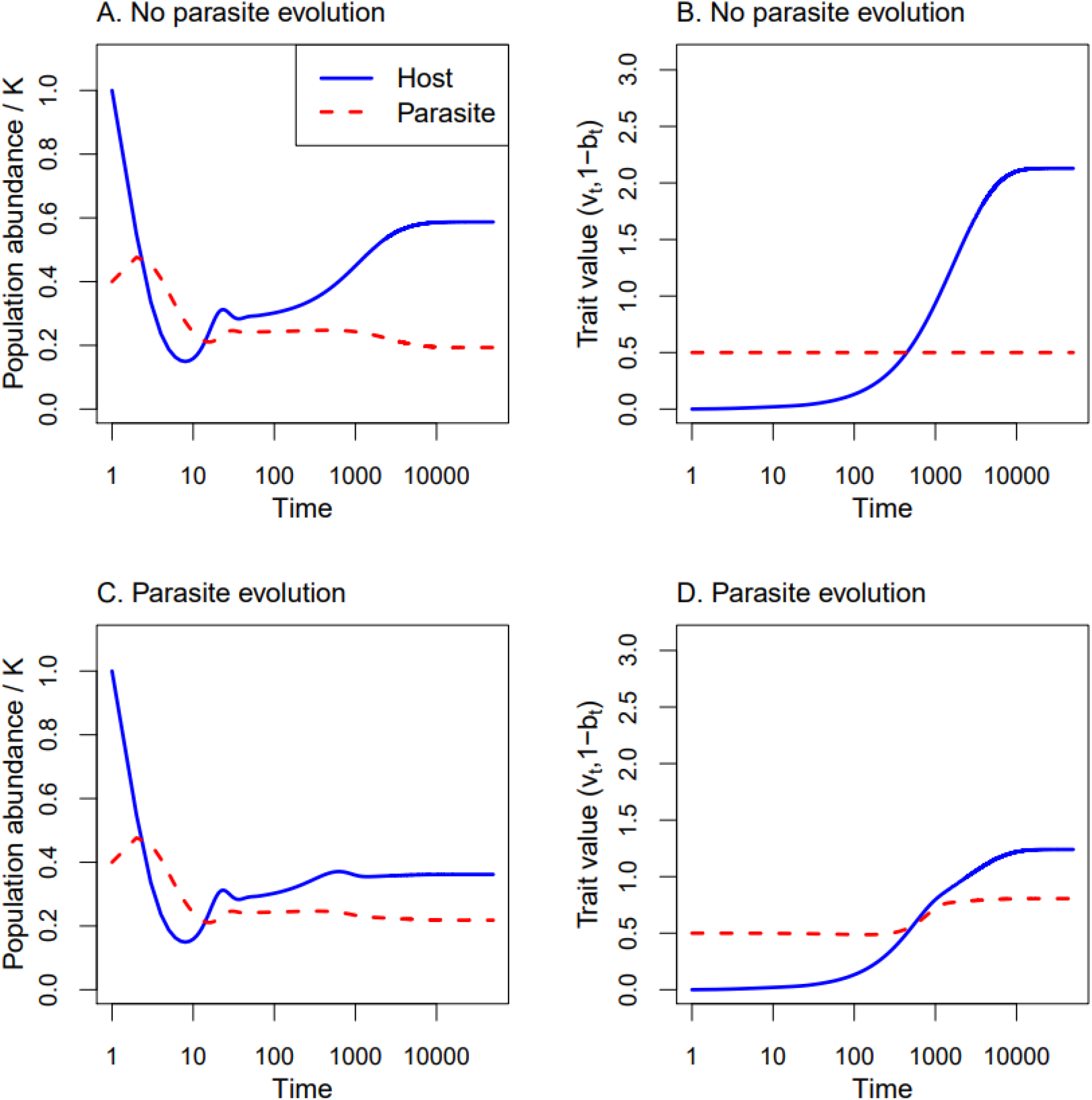
Time series of population abundances or parameter values for the host (blue) or parasite (red) populations. In (A),(B), there is no parasite evolution. In (C),(D), the parasite is allowed to evolve. Run with initial values *v*_0_=0.0001, *b*_0_=0.5, *N*_0_=0.5, *P*_0_=0.2, and parameters *K*=0.5, *r*_*n*_=0.5, *r*_*p*_=0.05, *q*=3, *f*=0.05, *g*=0.005, *m*=1.2, *c*=0.25, *V*_*N*_=0.01 for 50,000 time steps. For evolving parasite, *V*_*P*_=0.01, otherwise *V*_*P*_=0. Parasite trait values are 1 – *b*_*t*_, representing the parasite’s ability to overcome host defense, *v*_*t*_.

For some analyses, the equilibrium trait and population values of the runs were used. These were taken as the final values reached during the run, as observation found that by the 50,000 time steps used these values had long since stabilized.

Reintroductions of parasites can be modeled by overriding the parasite population *P* at regular time intervals. This was done by increasing *P* by its initial value *P*_0_. Any trait value changes between introductions were kept, to allow parasite evolution to proceed and determine where the equilibrium parasite trait would be in this scenario.

### Epidemiological model

In contrast to the resource consumer model where the parasite necessarily kills the host during transmission, the parasite/pathogen-host relationship can be represented through an epidemiological (SI) model (Anderson and May, 1979; Kermack et al., 1927). This approach has been used to consider spread and evolution of different parasite populations by tracking the susceptible (uninfected) population and separate populations of infection types, including those infected by the original strain, as well as a mutant strain (Levin and Pimentel, 1981; May and Anderson, 1983). We used a susceptible-infected model as described by May and Anderson (1983) as our model framework. The key differences are our inclusion of a clearance term, allowing infected individuals to return to susceptible at a constant rate. Unlike a typical susceptible-infected-resistant/recovered model (Anderson and May, 1979; Harko et al., 2014), we assume these recovered hosts can be infected again to reflect the lack of an adaptive immune system in arthropods (Rosales, 2017). We tracked host evolution instead by describing host ‘strains’: wild type and mutant. For simplicity, we assumed clonal reproduction without genetic recombination and did not model sexes separately. Thus, we had four populations, which were either susceptible *S* or infected *I*, and wild type *N* o*r* mutant *R*. The value of T represents the entire population, unless used as a subscript, in which case *T* represents the sum of the populations that share the classifier (*N*_*T*_ = *S*_*N*_ + *I*_*N*_; *I*_*T*_ = *I*_*N*_ + *I*_*R*_). These populations change according to the following equations:

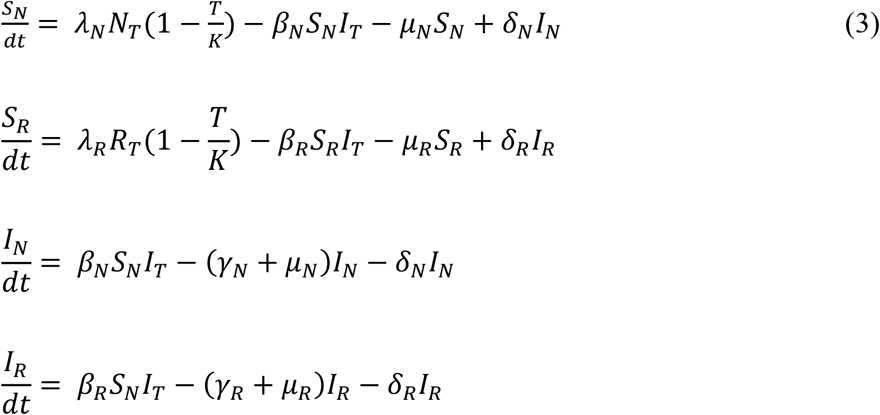

Here, *λ*_*i*_ is the per capita birth rate for host strain *i, μ*_*i*_ is its per capita mortality rate when uninfected, and δ_*i*_ is its per capita rate of clearance. The parameter β_*i*_ represents the pathogen’s transmissibility (rate of spread) for a given host strain, and *γ*_*i*_ its virulence (infection-caused mortality).

To simulate the models we started the mutant population as relatively small, and infection at wild type equilibrium levels, as though this mutation has just arisen and model its subsequent spread through the population. Both susceptible populations grow logistically, limited by the total population size with a carrying capacity of *K*. The mutant can differ from the wild type population in its transmissibility *β* and/or virulence *γ* parameters. This set difference in the effects of infection on the host populations assumes that any different effect of infection is entirely heritable, and that there is no environmental influence on phenotype.

We assume that the per capita birth rate *λ*_*R*_ is reduced in the mutant populations as a cost of the mutation, by setting *λ*_*R*_ < *λ*_*N*_. We have assumed infection of susceptible hosts is density dependent with respect to the sizes of the specific susceptible population and the total infected population, which assumes that infection can be spread to one host type from either host type and that the population is well mixed. This model distinguishes between two effective outcomes of host resistance – reduction of the biopesticide’s transmissibility, and reduction of the biopesticide’s virulence.

For considerations of manual infection augmentation, the equations had the addition of a term describing continuous environmental infection presumed to be through the application of the pathogen as a biopesticide. The infection occurs at a rate described by the per capita environmental infection rate *ζ*, which is multiplied by the susceptible population size. In the mutant case, this rate is scaled by the relative level of any transmission resistance the host has, _*T*_ assuming that resistance to horizontal transmission is similar to environmental resistance. The constant environmental infection rate allows for simplicity in the model, but assumes that the application rate of the pathogen is approximately equal to the pathogen degradation rate in the environment. Simulation for the model with environmental infection was the same as described above for the model without environmental infection. When including environmental infection, the equations were modified to be:

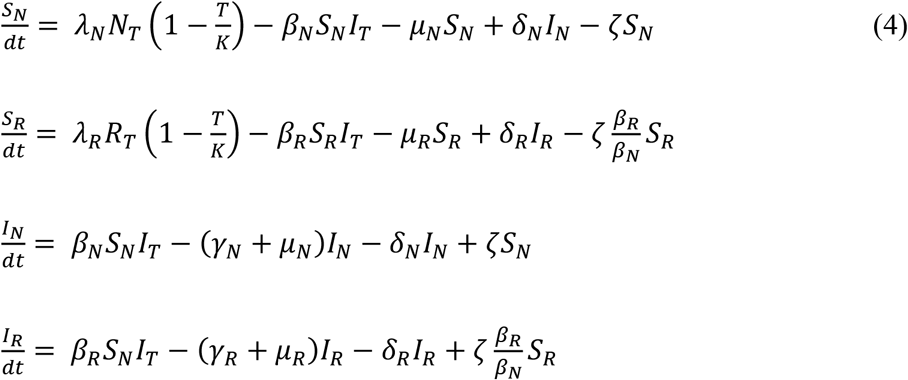

## Results and discussion

### Resource-consumer model

The first scenario we considered looked at the potential for biopesticide evolution in systems where the pesticide’s life cycle depends on its host’s mortality. This is the case in many nematodes (Ciche et al., 2006), and in entomopathogenic fungi that are forcibly ejected from cadavers (Steinkraus, 2006). Here, when a standard non-evolving parasite is introduced, there is an initial population reduction but the pest then evolves resistance and diminishes the amount of control provided (Figure 1A,B). When parasite evolution is allowed, however, pest population recovers much less and the pest evolves a lower level of resistance (Figure 1C,D). Since increasing resistance has associated costs, the final equilibrium density and trait value reached by the co-evolving host depends on the value of the resistance cost parameter, *f* (data not shown). Only a single stable state was observed from parasite evolution, so the parasite’s evolution would drive to its equilibrium value whether that meant increasing or decreasing its trait investment. In natural systems where there are multiple pathogens and hosts, governed by multiple traits previous studies suggest multiple stable states and complex dynamics (Northfield et al., 2021).

However, in the simpler case provided here, we can use the simple pairwise interaction to further evaluate the impacts of particular aspects of the system on the level of control at equilibrium. To better consider how biopesticide effectiveness might vary with the parasite trait in the absence of parasite evolution, we varied parasite trait value and evaluated the effects on the system’s equilibrium (Figure 2). At low parasite investment levels, increases in the parasite’s trait cause a reduction in the pest’s equilibrium population, but an increase in the equilibrium resistance investment by the pest. With increasing parasite strength however, eventually the trade-offs faced by the pest become too costly, and it can no longer sustain such high resistance investment. Its equilibrium resistance investment then decreases with increasing parasite trait value. There is an intermediate parasitic trait value in these results where both pest density and resistance were lowest. This trait value balances improving parasitic attack rates, with parasite growth. For parasite trait values more capable of overcoming prey defenses than this intermediate trait value, it is no longer beneficial to the pest to invest in defense, and thus there is no longer any benefit to the parasite of increasing trait investment to overcome host defense. Thus, investing further leads only to an increase in its trait costs and lower parasite density. In this case of high parasite trait levels, the parasite is unable to maintain a substantial population, allowing the pest population to grow. In sum, this analysis reveals that there is a parasite trait level that may provide the best pest control and improve pest resistance management. The equilibrium trait reached by the evolving parasite is quite similar to this value, but the best trait from the parasite’s perspective seems to be one that still allows some defense investment in its host. Importantly, these model analyses shows that biopesticide evolution can drive selection in the host toward minimal resistance investment, and minimal population size.

**Figure 2:**
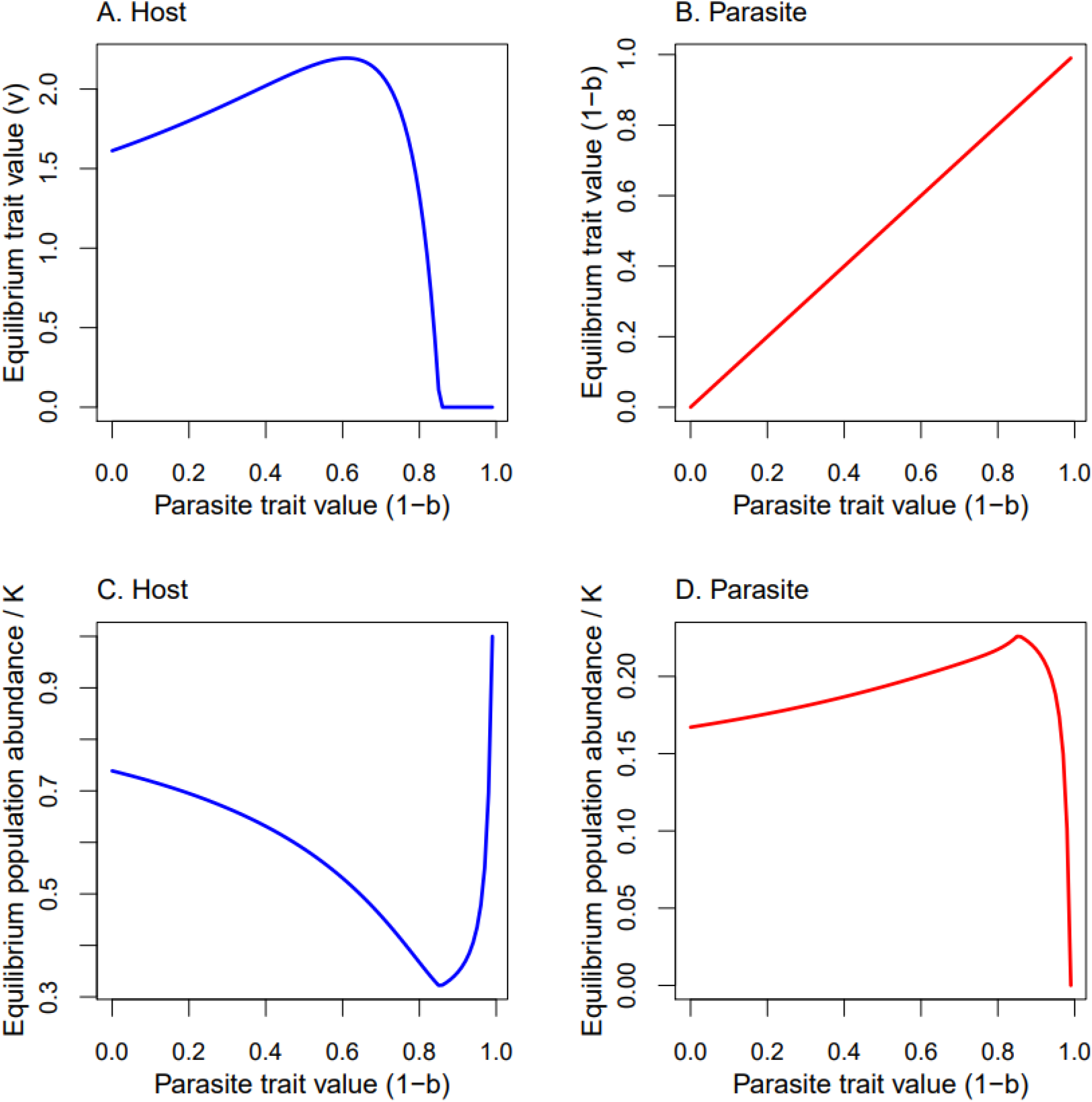
Equilibrium values of population abundances or parameter values for host (blue) and parasite (red) populations for various parasite parameter values, when parasite is not allowed to evolve. Run with initial values *v*_0_=0.0001, *N*_0_=0.5, *P*_0_=0.2, and parameters *K*=0.5, *r*_*n*_=0.5, *r*_*p*_=0.05, *q*=3, *f*=0.05, *g*=0.005, *m*=1.2, *c*=0.25, *V*_*N*_=0.01, *V*_*P*_=0, for 50,000 time steps. Parasite trait values are 1 – *b*, representing the parasite’s ability to overcome host defense, *v*.

A key difference between natural systems where host-parasite coevolution is often studied and pest management systems, is that in pest management the parasite densities are typically altered through repeated applications. Here, we simulated this by regularly supplementing the parasite population, to reflect regular parasite applications. We found that though this addition allows for slightly stronger parasites to persist despite their higher trait costs, the overall trend remains the same (Figure 3). In contrast to the single introduction, however, the traits have greater pest resistance investment compared to the single introduction, with slightly lower pest densities.

**Figure 3:**
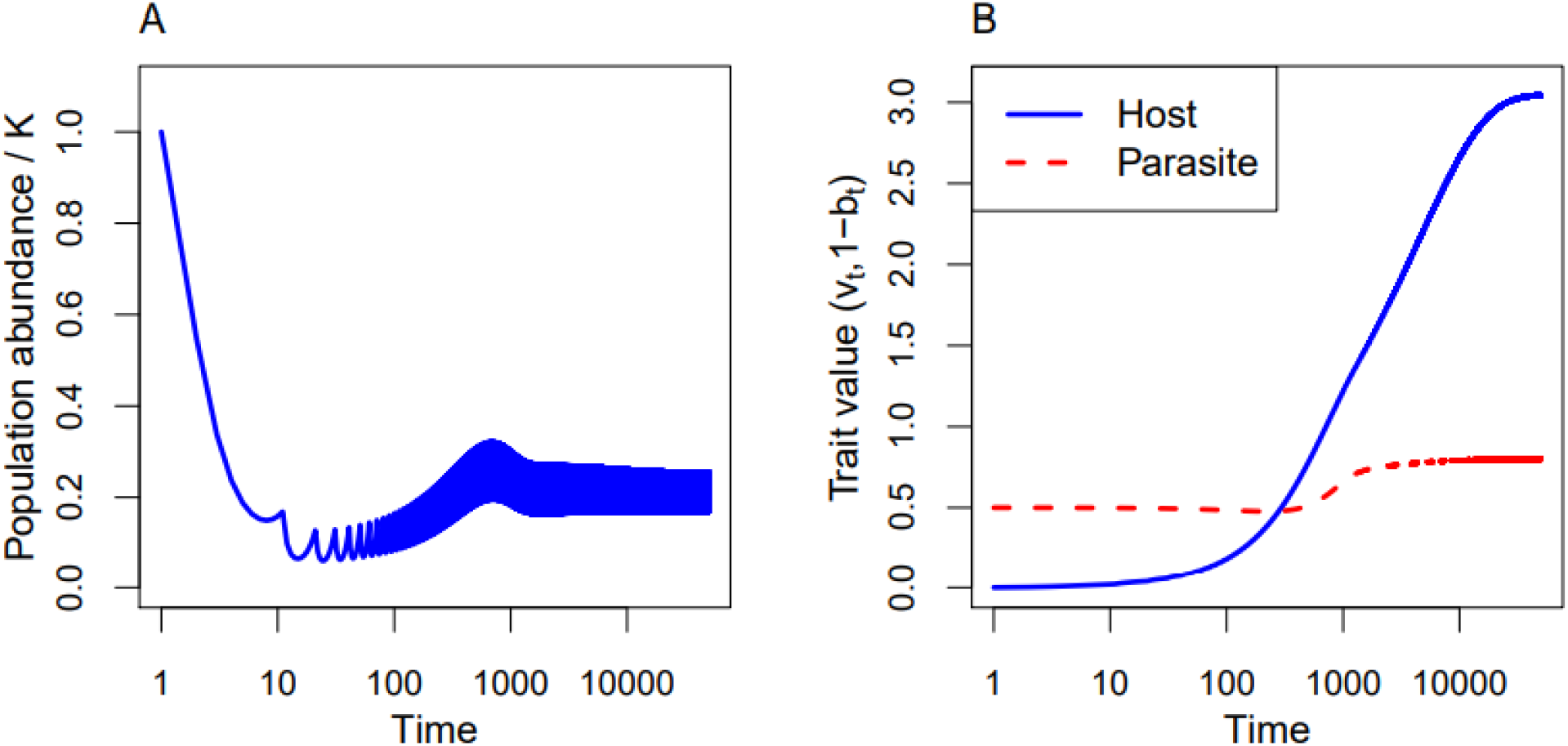
Time series of population abundances or parameter values for the host (blue) or parasite (red) populations with parasite evolution. Regular reintroductions of parasite population occur every 10 time steps, setting population *P* back to *P*_0_. Run with initial values *v*_0_=0.0001, *b*_0_=0.5, *N*_0_=0.5, *P*_0_=0.2, and parameters *K*=0.5, *r*_*n*_=0.5, *r*_*p*_=0.05, *q*=3, *f*=0.05, *g*=0.005, *m*=1.2, *c*=0.25, *V*_*N*_=*V*_*P*_=0.01 for 50,000 time steps. Parasite trait values are 1 – *b*_*t*_, representing the parasite’s ability to overcome host defense, *v*_*t*_.

### Epidemiological model

The second scenario considers systems more typical of infectious microbial pathogens, where a pathogen can be spread between living hosts. The benefits of different types of host defenses to disease, constitutive (reduced transmission) and induced (increased recovery) have been considered previously (Boots and Best, 2018). Here, we allow a beneficial mutation in the host to take the form of pathogen-tolerance or pathogen-resistance (Miller and Cotter, 2018). A pathogen-resistant host is one that is better able to prevent infection, as in codling moths’ type I resistance to granulovirus, where the resistance provides a systemic, early block on viral replication (Sauer et al., 2021). When we introduce a pathogen-resistant mutant to the wild type population, the overall infection prevalence (*I*_*N*_ *+ I*_*R*_) drops dramatically (Figure 4B). However, by reducing the total infection prevalence, this mutant indirectly decreases its own competitive advantage over the wild type, and the wild type hosts are able to continue to persist alongside them (Figure 4A). When considering varying degrees of both pathogen-resistance and pathogen-tolerance investment, the trend also holds that mutants with more pathogen-resistance suppress infection (Figure 5B,D) and support wild type persistence (Figure 5A). These findings support previous analyses of similar models showing that pathogen-resistant hosts will not reach fixation due to this phenomenon, and the pathogen will not be completely eradicated only by pathogen-resistance evolution (Roy and Kirchner, 2000).

**Figure 4:**
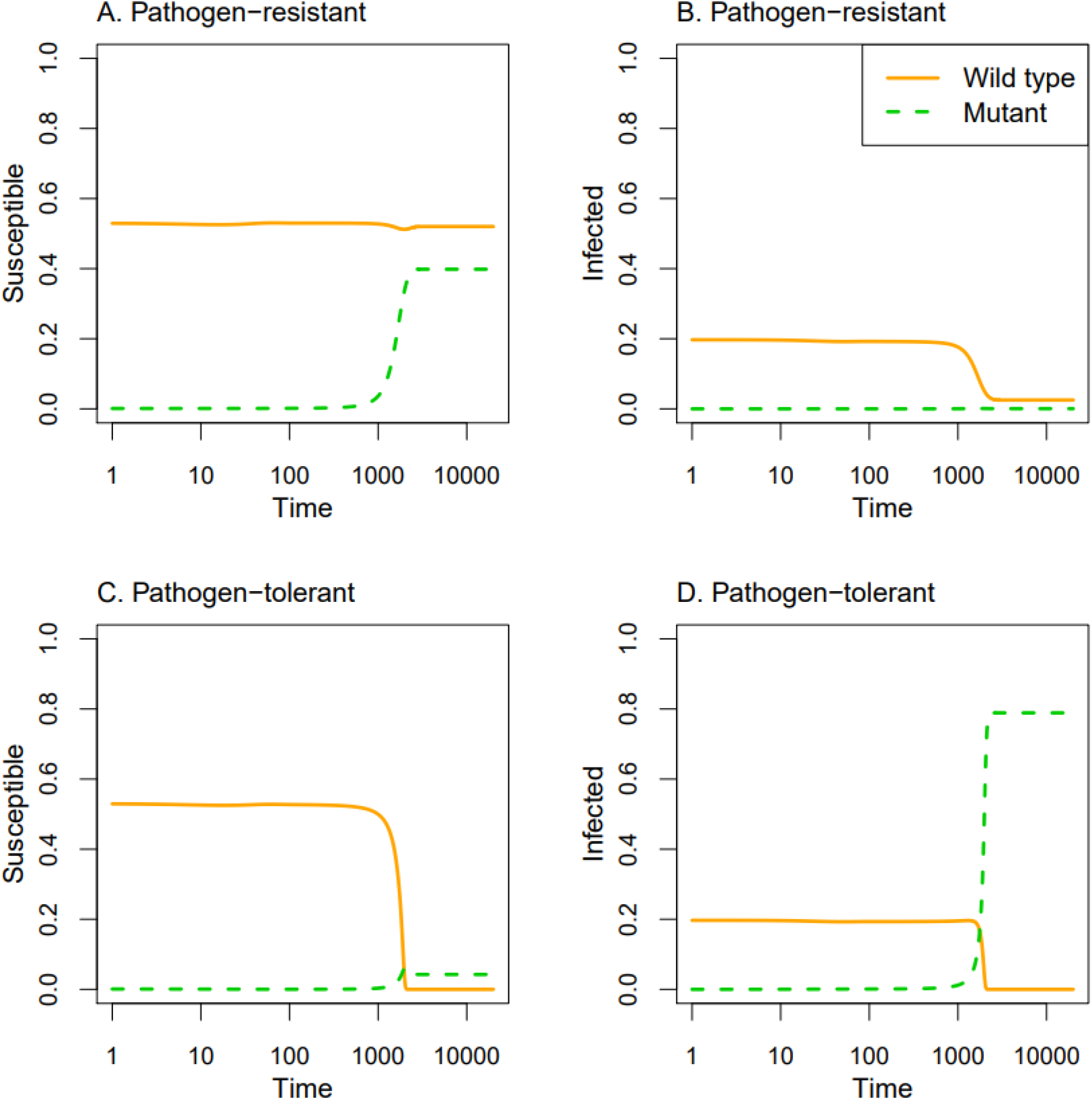
Time series of susceptible or infected population abundances for wild type (orange) and mutant (green) populations. In (A),(B), mutant experiences pathogen-resistance. In (C),(D), mutant experiences pathogen-tolerance. Run with initial values *S*_*N*_=0.530, *S*_*R*_=0.001, *I*_*N*_=0.197, *I*_*R*_ =0, and parameters 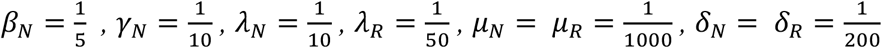 *K*=1 for 20,000 time steps. For pathogen-resistant mutant 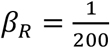, and for pathogen-tolerant mutant 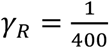. Otherwise *β*_*N*_=*β*_*R*_ and, *γ*_*N*_ *=, γ*_*R*_.

**Figure 5:**
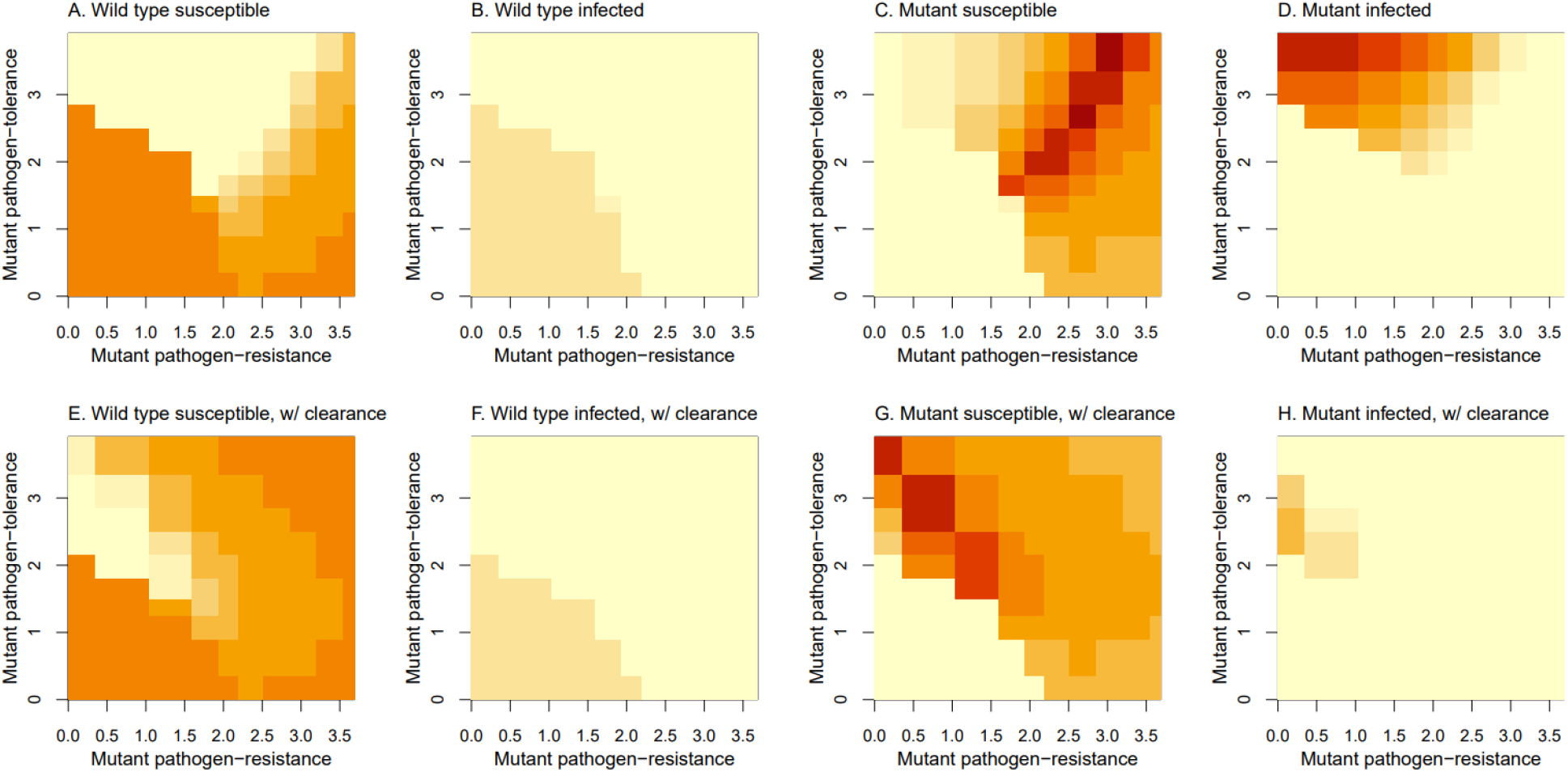
Equilibrium population abundances after introduction of a mutant for (A),(E) wild type susceptible, (B),(F) wild type infected, (C),(G) mutant susceptible, and (D),(H) mutant infected, at various levels of mutant pathogen-resistance and pathogen-tolerance, with darker colors signifying greater abundances and lighter colors signifying smaller abundances. Axes values are the log of the ratio between wild type and mutant traits, 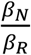 or 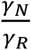. In (E - H) pathogen-tolerance is tied to a greater clearance rate. Run with initial values *S*_*N*_=0.998, *S*_*R*_=0.001, *I*_*N*_=0.001, *I*_*R*_=0, and parameters 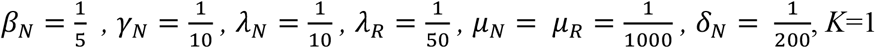, for 20,000 time steps. *β*_*R*_, *γ*_*R*_ vary from *β*_*N*_, *γ*_*N*_ to 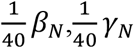. For (A-D) *δ*_*R*_ =*δ*_*N*_, and for (E-H), 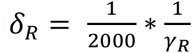.

A pathogen-tolerant mutant host that is better able to survive infection and is able to clear infection more readily can further be distinguished from one that simply survives and remains infected. This can be seen in burying beetles that can reduce entomopathogenic bacteria-related mortality rates by improving diet (Miller and Cotter, 2018). Here, we find that when pathogen-tolerance is conferred without improved clearance, the mutant can fully replace the wild type in its population (Figure 4C). The mutants essentially become reservoirs of infection and increase the overall infection in the system (Figure 4D). This then kicks off a positive feedback loop that promotes wild type displacement, where further increases in infection rates promote the selective advantage of the mutant strains. With the consideration of varying degrees of both pathogen-resistance and pathogen-tolerance investment, it again holds that more pathogen-tolerance promotes greater infection (Figure 5B,D) through the mutant population primarily comprising infected individuals (Figure 5C,D), and eliminates the wild types (Figure 5A,B). Previous analyses of similar models have shown that pathogen-tolerance should inevitably reach fixation when it arises (Roy and Kirchner, 2000). Roy and Kirchner (2000) also found this trend supported empirically in plant hosts of rust fungi, where the trend was for pathogen-resistance traits to be polymorphic and pathogen-tolerance traits to be fixed.

The positive feedback loop of increased pathogen-tolerance driving increased infection and fueling wild type displacement breaks down when pathogen-tolerance is linked with clearance of the pathogen from the host. When the mutant survives infection through recovery, the situation is similar to the pathogen-resistant case. When considering a range of both pathogen-tolerance and pathogen-resistance values, there are now more trait levels that can support wild type persistence (Figure 5E,F) and suppress infection (Figure 5F,H), with mutant populations now being primarily susceptible in most cases (Figure 5G,H). Here, the recovery can work to remove infection from the population, allow wild type persistence, and pressure the pathogen to evolve. Thus, pathogen-tolerance allows better wild type persistence when it is linked to clearance of the pathogen within the host, presumably through an effective immune response, as long as recovery has low levels of immunopathology (Cressler et al., 2015). For example, in cases where pathogen-tolerance protects against immunopathology (Medzhitov et al., 2012), there could then be a more sustained response improving recovery.

Since the potential for pathogen evolution to work as a strategy to improve biopesticide effectiveness thus depends on the effective type of mutation the host has, it is important to consider how the specifics of a given pest system impact the evolution of host and pathogen within artificial rearing environments. In the host-parasite scenario described above where transmission is necessarily linked to host mortality, our model analyses suggest that introducing pathogen-resistant hosts to the parasite rearing process should naturally select for parasites adapted to overcome host defense. This in theory would be a path forward to selecting for an improved biopesticide specifically “designed” to overcome pesticide resistance. However, the epidemiological model suggests the results may not be as straightforward when transmission regularly occurs between living hosts.

### Impacts on biopesticide resistance management

When the mutation is primarily a reduction of transmission (pathogen-resistance), our analyses suggest biopesticide evolution can select for stronger biopesticides. However, when the mutation is primarily a reduction of virulence (pathogen-tolerance), mutant populations can become fixed with overall infection much higher than prior to the mutation. Because the high pathogen-tolerance promotes pathogen prevalence and transmission (Roy and Kirchner, 2000), we do not expect pathogen evolution to lead to increased virulence that would improve biopesticide effectiveness.

Because pathogen-tolerance only improves host fitness when hosts are infected, and pathogen-resistance only improves host fitness for susceptible hosts in the presence of other infected hosts, the relative selection for each type of mutant (pathogen-tolerant or pathogen-resistant) will vary dynamically over the course of an epidemic and with the characteristics of the wild type and pathogen. Since these growth rates vary depending on the wild type’s parameters of infection (transmissibility, virulence), we compared the relative growth rates of each type of mutant in similar model simulations. In these scenarios, we treated pathogen-tolerant and pathogen-resistant mutants as experiencing a set fraction of the wild type’s transmissibility or Virulence 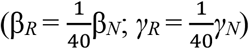.

Here, we found that more transmissible pathogens favored the evolution of pathogen-tolerant hosts while more virulent pathogens favored the evolution of pathogen-resistant hosts (Table 1). When the wild type experiences particularly high virulence (relative to transmission, e.g., *γ*_*N*_ > 1/8 when β_*N*_ =1/2 in our simulations) from the pathogen, mortality rates for infected individuals are still sufficiently large even if they have a reduction in virulence, so preventing infection is the better strategy. Likewise, when transmission is high (e.g., β_*N*_ > 1/2 when *γ*_*N*_ =1/8 in our simulations) susceptible hosts are still likely to contract infection even if their transmission is reduced, so being able to cope with infection is better (Table 1). These results align with previous analyses considering the conditions favoring two other manifestations of pathogen-resistance and pathogen-tolerance (Restif and Koella, 2004).

**Table 1:** Results of multiple model simulations comparing the different mutant host types (pathogen-resistant or pathogen-tolerant) to determine which has an earlier peak in population growth, without environmental infection. The cost of resistance was set as a 1/5 reduction in birth rate (high cost: 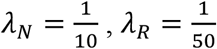, with 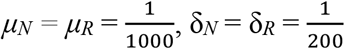, and *K*=1. In pathogen-resistant 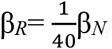, and in pathogen-tolerant 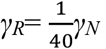; otherwise, β_*R*_= β_*N*_ and *γ*_*R*_= *γ*_*N*_. Other parameter values that were varied are presented in table column and row titles.

|  | **Virulence** |  |  |  |  |
| --- | --- | --- | --- | --- | --- |
| **Transmission** | | **Very Low**<br>($\gamma_N=1/20$) | **Low**<br>($\gamma_N=1/10$) | **High**<br>($\gamma_N=1/8$) | **Very High**<br>($\gamma_N=1/4$) |
| | **Very Low**<br>($\beta_N=1/10$) | Roughly<br>equivalent | Neither | Neither | Neither |
| | **Low**<br>($\beta_N=1/5$) | Tolerant | Resistant | Resistant | Neither |
| | **High**<br>($\beta_N=1/2$) | Tolerant | Tolerant | Roughly<br>equivalent | Resistant |
| | **Very High**<br>($\beta_N=3/4$) | Tolerant | Tolerant | Tolerant | Resistant |

To evaluate the infectivity for the different mutants, we also evaluated R_0_, the basic reproductive rate of the pathogen, (a measure of expected secondary infections from one individual in a susceptible population) for these different scenarios. May and Anderson (1983) gave the formula for R_0_ for this model, using variables consistent with this paper, R_0_ = β*T* / (*γ* + *μ* + δ). Here, R_0_ is proportional to transmission since it determines the amount of infected population growth, while virulence, background mortality, and clearance are inversely proportional since they determine the amount of infected population lost. Since R_0_ can be a function of a pathogen’s transmission and virulence, increasing with more transmission and decreasing with more virulence, this becomes a case of high R_0_ pathogens favoring pathogen-tolerance and low R_0_ pathogens favoring pathogen-resistance evolution in their hosts (Tables 1, 2). Again, this aligns with previous analyses (Restif and Koella, 2004).

**Table 2:** R_0_ values of the pathogen with respect to the wild type host at various pathogen virulence and transmission levels. R_0_ computed as R_0_ = *βT* / *(γ + µ + δ*), with 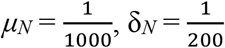 and T=1. Other parameter values that were varied are presented in table column and row titles.

|  | **Virulence** |  |  |  |  |
| --- | --- | --- | --- | --- | --- |
| **Transmission** | | **Very Low**<br>($\gamma_N=1/20$) | **Low**<br>($\gamma_N=1/10$) | **High**<br>($\gamma_N=1/8$) | **Very High**<br>($\gamma_N=1/4$) |
| | **Very Low**<br>($\beta_N=1/10$) | 1.786 | 0.943 | 0.763 | 0.391 |
| | **Low**<br>($\beta_N=1/5$) | 3.571 | 1.887 | 1.527 | 0.781 |
| | **High**<br>($\beta_N=1/2$) | 8.929 | 4.717 | 3.817 | 1.953 |
| | **Very High**<br>($\beta_N=3/4$) | 13.393 | 7.075 | 5.735 | 2.930 |

### Incorporating environmental infection

Not all virulence and transmission combinations can support a stable infection. This is the assumption in biopesticide scenarios, informing the repeated application of infectious agents. Low transmission reduces a pathogen’s ability to spread, while high virulence depletes the population of hosts that are able to spread it. At these levels, infection does not persist, so there is no selection pressure for hosts to evolve pathogen-tolerance or pathogen-resistance. Since R_0_ measures reproductive rate, an R_0_ less than 1 indicates that a pathogen population will not grow, and these levels of pathogen traits (Table 2) correspond with the simulation analyses (Table 1). Forcing infection to persist however, in a way that might reflect pathogen application as a biopesticide through regular augmentation of the infected population(s) independent of current infection size shows that the trends remain similar, with high virulence and low transmission scenarios that previously had no mutant growth now favoring pathogen-resistant mutants, with some shifting of where the switch between preferred mutation happens on the transmission/virulence axes (Table 3).

**Table 3:** Results of multiple model simulations comparing the different mutant host types (pathogen-resistant or pathogen-tolerant) to determine which has an earlier peak in population growth when environmental infection through pathogen application occurs. The cost of resistance was set as a 1/5 reduction in birth rate (high cost: 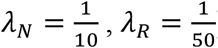, with 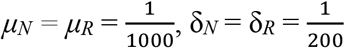, and *K*=1, and the application rate and effectiveness term was set to *ζ*=1/20. In pathogen-resistant 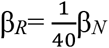, and in pathogen-tolerant 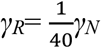, otherwise, β_*R*_= β_*N*_ and *γ*_*R*_= *γ*_*N*_. Other parameter values that were varied are presented in table column and row titles.

|  | Virulence |  |  |  |  |
| --- | --- | --- | --- | --- | --- |
| Transmission | | Very Low<br>($\gamma_N=1/20$) | Low<br>($\gamma_N=1/10$) | High<br>($\gamma_N=1/8$) | Very High<br>($\gamma_N=1/4$) |
| | Very Low<br>($\beta_N=1/10$) | Tolerant | Roughly equivalent | Resistant | Resistant |
| | Low<br>($\beta_N=1/5$) | Tolerant | Tolerant | Roughly equivalent | Resistant |
| | High<br>($\beta_N=1/2$) | Tolerant | Tolerant | Tolerant | Resistant |
| | Very High<br>($\beta_N=3/4$) | Tolerant | Tolerant | Tolerant | Resistant |

### Trait costs

In natural systems, many resistances do present additional costs to the mutant host’s fitness (Bartlett et al., 2018). It is further possible to quantify these costs: in another moth-granulovirus system, fitness costs of resistance were able to be calculated (Boots and Begon, 1993). These evolutionary trade-offs can be complicated though, and not directly linked to resistance genes (Bartlett et al., 2020). Here though, we considered simple (constitutive) costs associated with reproduction. To better see how one mutant strain would win in a scenario, this cost of mutation was allowed to vary. At increased mutation costs, several scenarios resulted in only one of the mutant types being able to invade. The trends for these aligned with those comparing earlier population growth, with high virulence favoring pathogen-resistant and high transmission favoring pathogen-tolerant hosts. Being able to predict which outcome of host evolution occurs would be beneficial as it would inform what options down the line should be available to deal with pest resistance as it develops in that system, and allow for costs associated with those management options to be considered when determining the optimal pest control strategy to implement. When considering a particular system, analysis of how host beneficial mutation costs are expressed and their impact on host fitness should allow for a more precise determination to be made.

Overall, when considering the epidemiological scenario, the decrease in pathogen presence shows an incentive for the pathogen to overcome its host’s pathogen-resistance. From this, the potential for pathogen evolution to produce a more effective biopesticide. In the pathogen-tolerant case however, the increase in pathogen presence may limit the potential for pathogen evolution to improve biopesticide effectiveness, as with an increased presence the pathogen would not be facing much incentive to overcome its host’s pathogen-tolerance.

There may be avenues for overcoming pathogen-tolerance, which itself should not reduce a pathogen’s fitness and may actually benefit pathogens through higher infection prevalence. Since transmission and virulence of a pathogen are often linked to some degree, a pure reduction in virulence may be difficult for hosts to achieve. Further, even if a pathogen in a pathogen-tolerant host is just as effective as it is in a wild type host, pressures may still exist to promote evolution of increased virulence. Hosts are often infected by multiple pathogens simultaneously, and if a mutation occurs within a host, it is now competing with other genotypes in the same host. In these cases, higher virulence could be considered something that allows that pathogen to outcompete the lower virulence pathogens within a host, as previous models have assumed (May and Anderson, 1983).

## Conclusion

When transmission and virulence of a biopesticide are completely linked, as when a parasite requires host death for transmission, and similar to a predator-prey relationship, the ability of biopesticide evolution as a means of dealing with pest resistance shows promise. When transmission and virulence can vary separately however, as when infection can occur between living hosts, there is more to consider. Host mutations that reduce pathogen transmission negatively impact infection populations, and so incentives for pathogen evolution to increase transmission likely exists; in contrast, host mutations that reduce pathogen virulence positively impact infection populations, and so there may not be incentives for pathogen evolution to increase virulence. This would mean that with pathogen-resistant hosts, biopesticide evolution could work as a management strategy, whereas with pathogen-tolerant hosts it would not. Real systems will likely fall somewhere in between these extremes of virulence-transmission linkage and beneficial mutation type, depending on the species involved and the environmental conditions where infection is occurring. Considerations of the outlook in these extremes though should help inform potential management strategies.

## Acknowledgements

R. Gomulkiewicz and D.W. Crowder provided helpful comments on an earlier draft of this manuscript.

## Funding

Funding was provided by a BIOAg grant from Washington State University.

## Notes

### Competing Interest Statement

The authors have declared no competing interest.

